# Cumulative effects of mutation and selection on susceptibility to bacterial pathogens in *Caenorhabditis elegans*

**DOI:** 10.1101/2021.09.07.459309

**Authors:** Sayran Saber, Lindsay M. Johnson, Md. Monjurul Islam Rifat, Sydney Rouse, Charles F. Baer

**Author notes:** Communication to. These authors contributed equally (Sayran Saber, Lindsay M. Johnson).

## Abstract

Understanding the evolutionary and genetic underpinnings of susceptibility to pathogens is of fundamental importance across a wide swathe of biology. Much theoretical and empirical effort has focused on genetic variants of large effect, but pathogen susceptibility often appears to be a polygenic complex trait. Here we investigate the quantitative genetics of survival over 120 hours of exposure ("susceptibility") of *C. elegans* to three bacterial pathogens of varying virulence, along with the standard laboratory food, the OP50 strain of *E. coli*. We compare the genetic (co)variance input by spontaneous mutations accumulated under minimal selection to the standing genetic (co)variance in a set of 47 wild isolates. Three conclusions emerge. First, mutations increase susceptibility to pathogens, and susceptibility is uncorrelated with fitness in the absence of pathogens. Second, the orientation in trait space of the heritable (co)variance of wild isolates is sufficiently explained by mutation. However, with the possible exception of *S. aureus*, pathogen susceptibility is clearly under purifying, directional, selection of magnitude roughly similar to that of competitive fitness in the MA conditions. The results provide no evidence for fitness tradeoffs between pathogen susceptibility and fitness in the absence of pathogens.

## Introduction

The relationship between pathogens and their hosts holds an exalted place in evolutionary biology (Haldane 1949; Hamilton 1980). It is well-established that susceptibility of individuals to the harmful effects of pathogens often has a genetic basis. The notion that *particular* multilocus host genotypes confer resistance or susceptibility to *particular* pathogen genotypes has been especially important, because "Red Queen" models of the evolution of sex and recombination are predicated on fluctuating epistasis (Barton 1995), with negative frequency-dependent selection resulting from a host-pathogen arms-race providing the most obvious plausible scenario (Hamilton 1980).

In some cases, the genetic basis of variation in host susceptibility to pathogens can be attributed to variants at one or two loci. For example, variants at a few loci in the human genome (e.g., *HBB*, *ABO*, *G6PD*) are famously implicated in resistance to malaria (Rockett et al. 2014). Often, however, the heritable basis of susceptibility to a particular pathogen appears to be polygenic, with a substantial fraction of unexplained heritability (Lefebvre and Palloix 1996; Hill 2001; Daub et al. 2013; Cogni et al. 2016), sometimes even in cases with segregating large-effect loci (e.g., malaria; Band et al. 2019). Pathogens may thus impose significant selection even in the absence of Red Queen dynamics. It seems reasonable, then, to consider "susceptibility to pathogen X" as a generic complex trait; in some cases, there will be loci of large effect that explain much of the variance in susceptibility, but other times there will not.

Two fundamental issues in understanding the biology of complex traits, are (*i*) what is the rate of input of genetic variance (and covariance) by mutation? and (*ii*) what is the relationship of the trait(s) with fitness? Taken together, these two considerations present a hierarchy of evolutionary hypotheses. The simplest explanation for any pattern of (co)variation is neutral evolution *sensu stricto*, i.e., the pattern can be sufficiently explained by neutral mutation and random genetic drift. If the pattern cannot be plausibly explained by neutral evolution, the next-simplest explanation is purifying selection against unconditionally deleterious mutations, plus drift. Only if those two explanations prove unsatisfactory need more complicated explanations be considered; for example, scenarios of balancing or positive selection.

For quantitative traits, the analog of the per-nucleotide mutation rate, μ, is the mutational variance *V_M_*=*U*α^2^, where U_M_= ∑*_i_* µ*_i_* is the genomic mutation rate and α^2^ is the average squared effect of a new mutation on the trait (Clayton and Robertson 1955; Lynch and Hill 1986; Barton 1990). For a neutral trait at mutation-drift equilibrium, the genetic variation segregating within a population *V_G_*=2*N_e_V_M_*, and populations will diverge asymptotically at rate 2*V_M_* (Lynch and Hill 1986). For traits under purifying selection (directional or stabilizing), at mutation-selection balance (MSB), 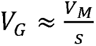, where *s* is the average strength of purifying selection against a mutant allele that affects the trait (Barton 1990; Kondrashov and Turelli 1992; Charlesworth 2015). The relationship 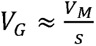 implicitly includes both the direct selective effects of mutations on the trait in question, but also the pleiotropic effects of selection on correlated traits.

The multivariate extensions of *V_G_*and *V_M_* are the standing genetic and mutational variance-covariance matrices, **G** and **M**, of which the diagonal elements are the variances, *V_G,i_* and *V_M,i_* respectively, and the off-diagonal elements are the genetic/mutational covariance between traits *i* and *j*, *Cov_G,ij_* and *Cov_M,ij_*(Lande 1980; Phillips and McGuigan 2006). The covariances capture the cumulative effects of pleiotropy and linkage disequilibrium.

Here we report results of a set of experiments in which we (*i*) estimate the cumulative effects of spontaneous mutations on the susceptibility of two strains of *C. elegans* to three bacterial pathogens, and (*ii*) estimate the standing genetic (co)variance for susceptibility to those same three pathogens from a set of ∼50 *C. elegans* wild isolates. Our first, broad goal is to determine the realms of consistency and of idiosyncrasy in the mutational process in the context of host-pathogen evolution – what features are specific to particular host genotypes, or particular pathogens, and what features are general. Our second, broad goal is to draw inferences about the strength and pattern of natural selection acting on *C. elegans*’ susceptibility to bacterial pathogens, relative to the inferred strength of selection on competitive fitness.

## Materials and Methods

### Mutation Accumulation (MA) experiments – basic principles

An MA experiment is simply a pedigree, in which replicate populations derived from a known, (ideally) isogenic progenitor are propagated under conditions in which natural selection is minimized (Mukai 1964; Halligan and Keightley 2009). Typically, selection is minimized by minimizing *N_e_*; mutations with fitness effects *s*<1/4*N_e_* are effectively neutral (Keightley and Caballero 1997). MA experiments can be used to illuminate the workings of natural selection in two ways. First, they provide the most straightforward and least assumption-laden way to estimate mutational (co)variances, which can be compared to the standing genetic (co)variances as outlined in the Introduction (**Supplemental Figure S1**). Second, if selection is directional, the change in the trait mean (the mutational bias, ΔM) will point in the opposite direction of selection. On average, fitness declines under MA conditions. Traits that are positively correlated with fitness should, on average, exhibit a negative mutational bias (ΔM<0); traits that are negatively correlated with fitness should, on average, exhibit a positive mutational bias (ΔM>0).

### Mutation Accumulation (MA) lines

Details of the mutation accumulation experiment are given in Baer et al. (2005). N2 is the standard laboratory strain of *C. elegans*; PB306 is a wild isolate graciously provided by Scott Baird. The basic protocol follows that of Vassilieva and Lynch (1999) and is depicted in **Supplemental Figure S2**. Briefly, replicate populations (MA lines) were initiated from a cryopreserved stock of a highly-inbred ancestor and propagated by transferring a single immature hermaphrodite at each generation (four-day intervals) under standard laboratory conditions. The lines were propagated for 250 transfers (G_250_) prior to cryopreservation. Data on transfers and additional context is given in **Supplemental Table S1**. Of the 100 N2 and PB306 MA lines initiated in 2001, we report results from 59 N2 and 65 PB306 lines. On average, each MA line carries about 65 spontaneous base-substitution and small indel mutations (Rajaei et al. 2021), with a few larger structural variants (Saxena and Baer, unpublished results). The average number of mutations per genome is nearly identical between the N2 and PB306 MA lines, thus any differences between the two strains in the cumulative effects of mutations are the result of differences in the distribution of mutational (and/or epimutational) effects, rather than in the total number of mutations.

The cryopreserved progenitor (G0) serves as a control in two ways. The difference between the trait mean of the G0 and MA treatments (ΔM) quantifies the cumulative effect of spontaneous mutations on the trait mean. If the G0 progenitor is subdivided into replicate "pseudolines" (PS) which are subsequently treated identically to the MA lines (**Supplemental Figure S2**), the difference between the among-MA line and the among-PS line components of variance represents the cumulative heritable variance resulting from the accumulation of spontaneous mutations (and potentially epimutations, depending on the experimental design; Beltran et al. 2020; Johnson et al. 2020). PS lines were constructed by thawing a sample of the cryopreserved G0 progenitor and picking individual L4 stage hermaphrodites to new plates, each of which was designated a PS line. PS lines were subsequently treated as if they were MA lines. Unfortunately, due to a management error on the part of the senior author (CFB), not all pathogen treatments include similar numbers of pseudolines (see below).

### Competitive fitness assay

Competitive fitness of the MA lines and their G0 progenitors was assayed in May, 2005. The details of that assay have been published elsewhere (Yeh et al. 2018), and are reprised in section 4 of the **Supplemental Material**. Fitness data are given in **Supplemental Table S2**.

### Wild isolates

A collection of wild isolates of *C. elegans* was obtained from Erik Andersen (Northwestern University) and cryopreserved in the Baer lab. A list of the wild isolates is given in **Supplemental Table S3**. The genome sequences of the wild isolates along with collection information are available at https://caendr.org/.

### Pathogen infection and survival assay

We used four strains of bacteria; three documented pathogens (the PA14 strain of *Pseudomonas aeruginosa*, the NCTC8325 strain of *Staphylococcus aureus*, and the OG1RF strain of *Enterococcus faecalis*) and the standard laboratory worm food, the OP50 strain of *E. coli*. We use the term "pathogen" to refer to the three pathogenic species, and "bacteria" when OP50 is included, although there is reason to think that OP50 is a weak pathogen of *C. elegans*, on the grounds that lifespan is extended when worms are fed on killed bacteria compared to live bacteria (Portal-Celhay et al. 2012). For the MA lines, each bacterial species was assayed in three or four experimental blocks. Wild isolates were assayed in one assay block per species. The basic protocol is a variant of the "slow killing assay" (Etienne et al. 2015), in which ∼30 L4-stage worms are transferred to a plate spread with bacteria and worms scored as live or dead at 12-hr intervals over the course of 120 hrs. The assay is described in detail in section 1 of the **Supplemental Material**, and depicted in **Supplemental Figure S3**.

### Data Analysis

#### (i) Survival probability

Relative survival of a focal type (replicate *i* of strain *x* of treatment *y* on bacteria *z*) was quantified as the fraction of worms *f* (*f=n_LIVE_/n_TOTAL_*) surviving at time *t* relative to the mean fraction surviving of the N2 G0 ancestor on OP50, *f_REL_*= *f_FOCAL_* / *f_N2,OP50_*. For all bacteria except PA14, survival was quantified at the culmination of the assay, at *t*=120 hrs. Survival was much lower on PA14 than on the other strains (**Supplemental Figure S4**), so survival on PA14 was quantified after 60 hrs. Complete survival data from the MA assay are given in **Supplemental Table S4**. Times were chosen to roughly maximize the total variance in survival (**Supplemental Table S5**).

#### (ii) Mutational parameters

Two statistics were calculated to quantify the cumulative effects of mutation accumulation on pathogen susceptibility. First, we estimated the mutational bias (ΔM), the per-generation change in the trait mean; ΔM = (*f_REL,MA_-f_REL,G0_*)/*t*, where *t* is the number of generations of MA (*t*=250) (Lynch and Walsh 1998).

Second, we measured the mutational variance, V_M_, calculated as (V_L,MA_-V_L,0_)/2*t*, where V_L,MA_ is the among-line variance of the MA lines, V_L,0_ is the among-line variance of the PS lines, and *t* is the number of generations of MA. V_M_ standardized as a proportion of the residual (within-line) variance, V_E_, is the "mutational heritability";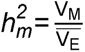. Mutational covariances, Cov_M_, were calculated analogously to V_M_, from the among-line components of covariance.

Genotypic variances (V_G_) and covariances (Cov_G_) of the wild isolates were calculated similarly to the mutational (co)variances. There is very little residual heterozygosity in the wild isolates (Rajaei et al. 2021), so wild isolates were treated as inbred lines, in which the genotypic variance V_G_ = V_L_ (Zhang et al. 2021). Genetic covariances and correlations were calculated analogously. Broad-sense heritability, 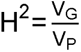, where V_P_ is the total phenotypic variance. We note that there is disagreement within the community of worm quantitative geneticists whether H^2^ should be estimated from V_L_ or V_L_/2. The former assumes that the base population is already inbred; the latter assumes that the base population is outbred (Falconer 1989). The traditional calculation of H^2^ from inbred lines assumes the latter (e.g., the popular software package H2boot; Arnold and Phillips 1999), but most worm quantitative geneticists use the former. For the sake of comparison with other studies, we have capitulated and use 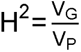 (note that Etienne et al. (2015) used the traditional V_L_/2 formulation in their calculation of H^2^).

### Statistical Analysis

To account for variation among assay blocks, we first applied the general linear model (GLM) *y_ij_*= μ*+b_j_+e_i_*_j_ to each combination of strain (N2 or PB306) and bacterial species independently, where *y_ij_* is the value of *f_REL_* of replicate *i* in assay block *j*, μ is the overall mean, *b_j_* is the fixed effect of block *j*, and *e_ij_* is the residual effect, assumed normally distributed with mean 0. The residuals of the linear model, *y\**, are the dependent variable in subsequent analyses. Pooled over strains, treatments (MA, G0) and bacteria, the *y\** are normally distributed (**Supplemental Figure S5**); we removed outliers with absolute Studentized residuals >3 (n=10/2021 samples).

Next, still considering each combination of *C. elegans* strain (N2 or PB306) and bacterial species independently, we estimated means and variances from the GLM *y*_ijk_=*μ*+a_i_+L_j|i_+e_jk|i_*, where *y*_ijk_* is the residual from the block mean, μ is the overall mean, *a_i_* is the fixed effect of generation of MA (gen=0 for MA, gen=250 for MA), *L_j|i_*is the random effect of line (MA or pseudoline) within each treatment group, and *e_jk|i_* is the residual effect, assumed normally distributed with mean 0. Variances of random effects were estimated independently for each treatment group (banded main diagonal covariance structure). Variances were estimated by restricted maximum likelihood (REML) with degrees of freedom determined by the method of Kenward and Roger (1997), as implemented in the MIXED procedure of SAS v.9.4. The fixed effect of treatment was tested by F-test with Type III sums of squares. Significance of among-line variance components (V_L_) was assessed by likelihood ratio test (LRT) of the model with the among-line component of variance included against the model with the among-line variance constrained to 0. Since V_M_ and 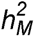 are composite parameters, confidence intervals were estimated by means of a parametric bootstrap, assuming that the components (V_L,MA_, V_L,PS_, V_E_) are normally distributed with means equal to the observed and standard deviations equal to the estimated standard errors. Individual bootstrap estimates of variances were constrained to be non-negative, i.e., bootstrap replicates less than 0 were set equal to 0.

Genetic covariance matrices (**M** and **G** for MA lines and wild isolates, respectively) were estimated analogously; the details are given in section 2 of the **Supplemental Material**.

The orientations of **M** and **G** in multivariate trait space were compared using the "genetic line of least resistance" approach, following Schluter (1996). We describe the method here; justification is provided in the Results. Data were standardized by the block standard deviation (i.e., Studentized residuals, *y’*), to put the four variables on a common scale. **M** and **G** were used as input data in a principal components analysis (PCA), as implemented in the PRINCOMP procedure in SAS v. 9.4. The first principal component (PC1) is the axis of maximum genetic variance, and evolution will proceed most rapidly and easily along this axis, be it due to selection or drift (Lande 1979; Phillips et al. 2001). PC1 of **G** is commonly referred to as **g**_max_ (Schluter 1996); we follow that convention and refer to PC1 of **M** as **m**_max_. The deviation in orientation in trait space between **m**_max_ and **g**_max_ is represented by the angle θ between the two vectors, where θ=cos^-1^(**m**_max_ ^T^**g**_max_) (Schluter 1996).

If **g**_max_ = **m**_max_, θ=0. We tested the hypothesis that **g**_max_ = **m**_max_ as follows. All MA lines (n=124, pooled over strains) and wild isolates (n=47) were pooled into one dataset, and line type (MA or WI) was assigned to each line randomly, with 124 random lines being designated "pseudoMA" and the remaining 47 lines designated "pseudoWI". We re-calculated the among-line (co)variance matrices and re-ran the PCA as described above; we refer to PC1 of the randomized data as **m**_max_**’ and g**_max_**’**, respectively, and the angle between them as θ*_rand_*. We repeated this procedure 1000 times; the fraction of replicates for which θ*_rand_*> θ is the approximate P-value of a test of the null hypothesis that **g**_max_ = **m**_max_.

## Results and Discussion

Our overarching purpose is comparative evolutionary genetics in the context of host-pathogen relationships – what features are specific to particular host genotypes, or particular pathogens, and what features are general. Survivorship of the N2 G0 progenitor on OP50 (food) sets the benchmark by which the effects of the pathogenic bacteria are assessed. On average, the N2 progenitor suffered approximately 11% mortality over the course of the 120 hr assay (note that the assay was conducted at 25°, which is stressfully warm for *C. elegans*). Relative to that benchmark, survivorship was further reduced by as little as 1.5% in the PB306 G0 progenitor on OP50 and by as much as 84% in the N2 G0 progenitor on PA14. Averaged over the two strains (N2 and PB306) and treatment groups (MA and G0), the relative pathogenicity of the three pathogenic bacteria is OP50<*E. faecalis*<*S. aureus*<<PA14 (**Supplemental Figure S5; Supplemental Table S6**). Interestingly, the relative pathogenicity was different for the wild isolates, for which the rank order of pathogenicity was *E. faecalis*<*S. aureus*<OP50<<PA14 (**Supplemental Figure S5**. The discrepancy appears to be due to N2 and PB306 having substantially higher than average survivorship on *E. coli* OP50. PB306 was included in the wild isolate assay and its relative survival was similar to the MA assay for all bacterial species except *E. faecalis* (albeit n=3 replicates in the WI assay vs. >100 replicates in the MA assay). N2 has been selected to eat OP50 for thousands of generations, so *E. coli* (or at least OP50) should probably be considered a pathogen (or at least as "unhealthy food") for all strains except N2. It is well-known that worms live longer on killed OP50 than on live OP50 (Portal-Celhay et al. 2012), so its status as a pathogen has some precedence.

### Mutation Accumulation (MA) lines

#### (i) Means

We expect that survival on OP50 (food, at least for N2) should be unambiguously beneficial, and both N2 and PB306 exhibit a significant negative mutational bias on OP50 (**Table 1**; **Figure 1A, B**). Predictions about the direction of selection for survivorship on pathogenic bacteria are less clear, because fitness tradeoffs between pathogen susceptibility and traits such as growth rate and yield in the absence of pathogens are well-documented (Biere and Antonovics 1996; Tian et al. 2003; Kraaijeveld and Godfray 2008; Duncan et al. 2011).

**Figure 1.**
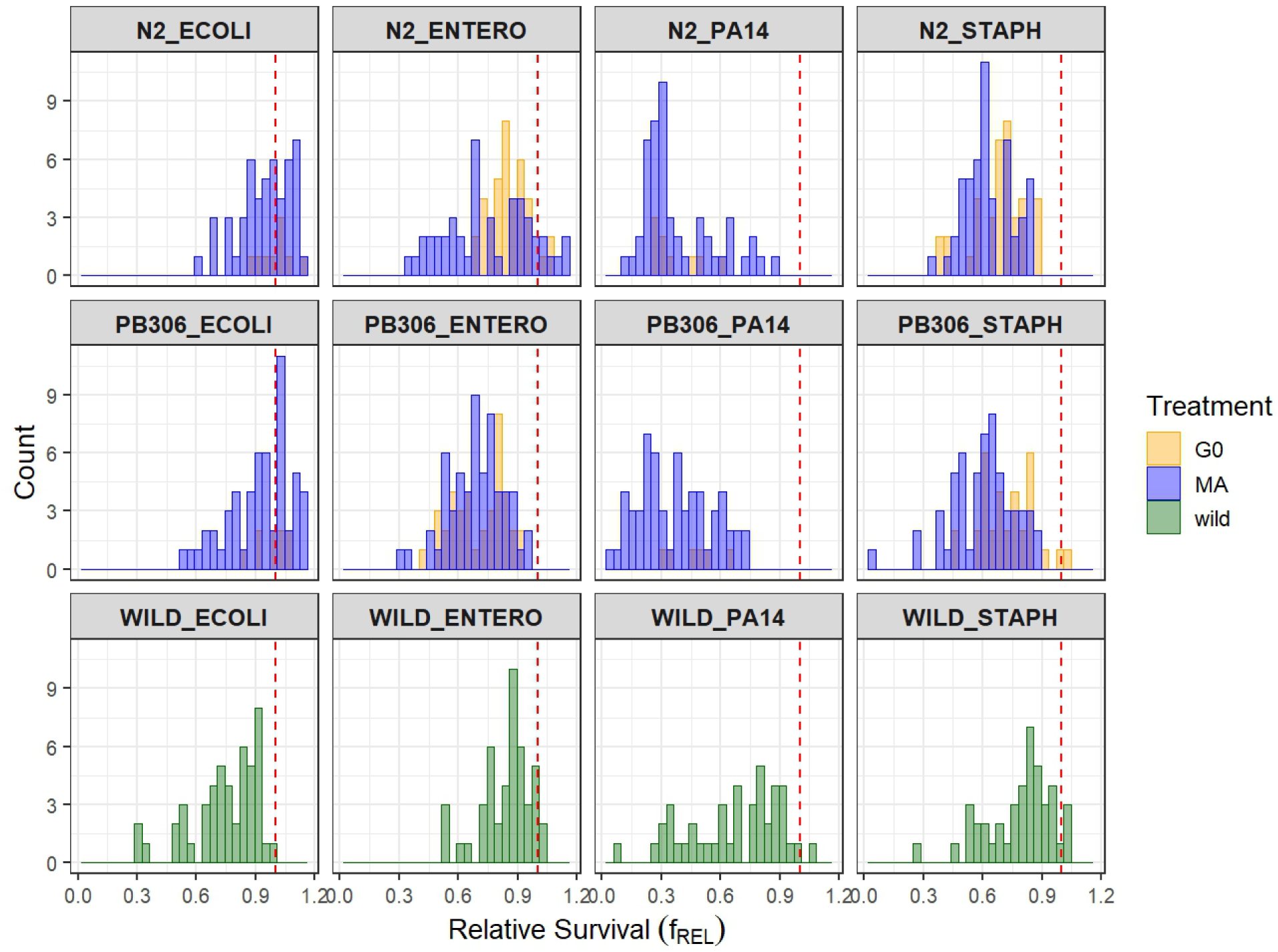
Distribution of line mean relative survival (*f_REL_*; see Methods for explanation) on the four bacterial species (left to right: *E. coli* OP50, *Enterococcus faecalis*, *Pseudomonas aeruginosa* PA14, *Staphylococcus aureus*). G0 MA ancestor in orange, MA lines in blue, Wild isolates in green. Red dashed line (*f_REL_*=1) is the mean survival of the N2 G0 lines on *E. coli* OP50. (A) Top panel; N2; (B) Middle panel, PB306; (C) Bottom panel, wild isolates.

**Table 1.**
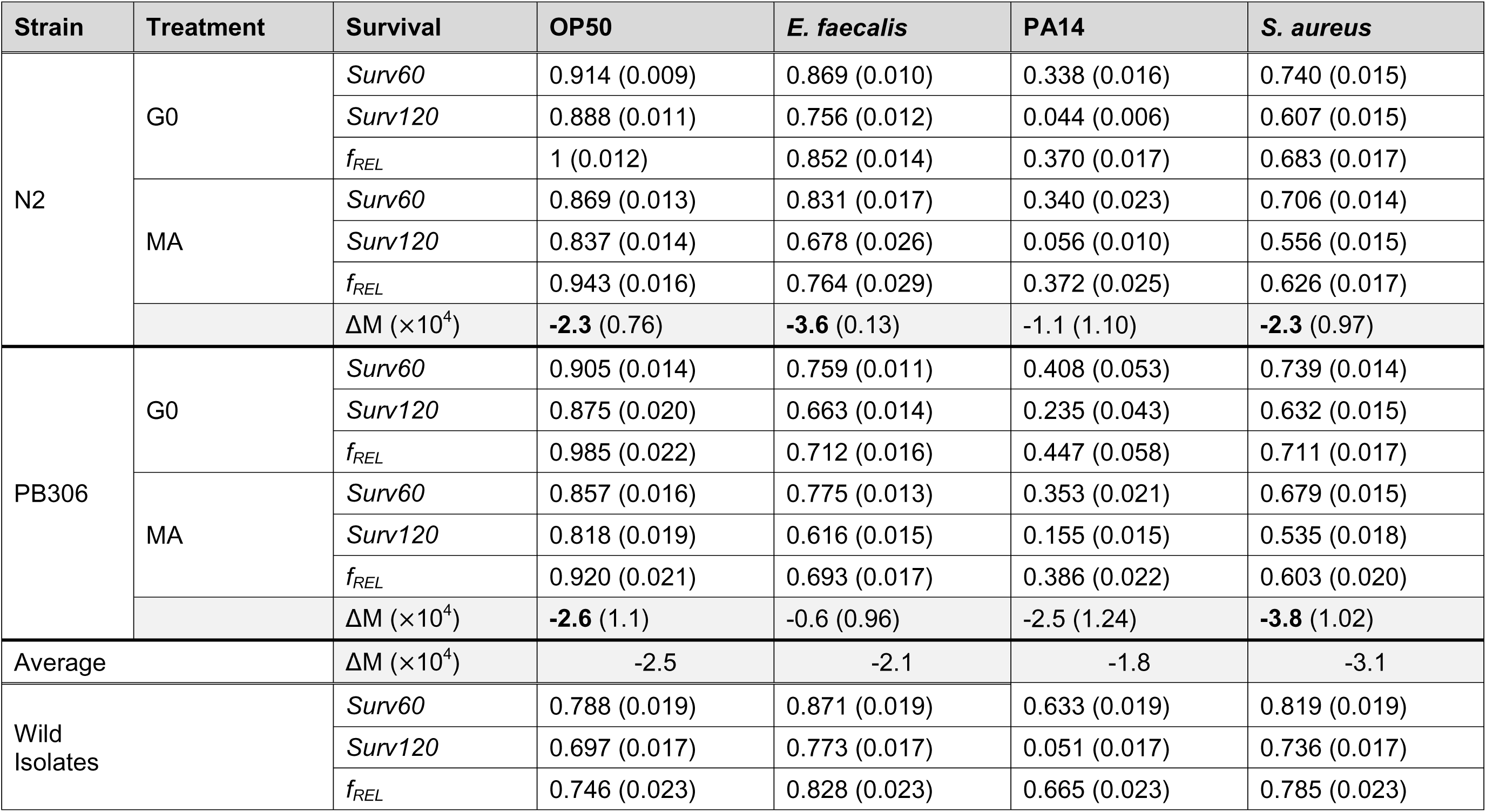
Means (SEM). Survival measures are: *Surv60*=% surviving at 60 hrs; *Surv120*=% surviving at 120 hrs; see Methods for details of calculations of *f_REL_* and ΔM. Values of ΔM in bold font are significantly different from 0 (P<0.05); see Methods for details of statistical tests.

Whether the deleterious effects of mutations are exacerbated under stressful conditions is a topic of active investigation (e.g., Matsuba et al. 2013; Yun and Agrawal 2014; Katju et al. 2018), but there is no theoretical reason to expect them to be unless selection is soft (Agrawal and Whitlock 2010), which it is not in these assays. Infection by a pathogen is *prima facie* stressful. If deleterious effects are greater under pathogen stress, we predict that ΔM will be greater (more negative) in the pathogen treatments than in the OP50 treatment, and that ΔM will be greatest (most negative) in the most virulent pathogen, PA14.

As a whole, support for those predictions is weak. Point estimates of ΔM are negative in all eight strain/bacteria combinations (**Table 1**), although not significantly different from zero in three of the eight, and less on PA14 than on OP50. In only two cases – N2 on *E. faecalis* and PB306 on *S. aureus* – is ΔM substantially greater (∼1.5X) in the pathogen than on OP50.

#### (ii) Variances

To begin, we estimated the among-line component of variance (V_L_) for each combination of bacteria, strain, and treatment (MA or G0). For the MA lines, V_L,MA_ includes the combined contributions of new mutations and heritable, non-genetic effects; for the G0 PS lines, V_L,PS_ includes only the heritable, non-genetic effects. V_L,MA_ is significantly greater than zero in five of the eight strain/pathogen combinations in the MA lines, and nearly significant in a sixth (PB306 in *E. faecalis*; LRT p<0.07). There was no significant V_L,MA_ on *S. aureus* in either strain (**Table 2**).

**Table 2.**
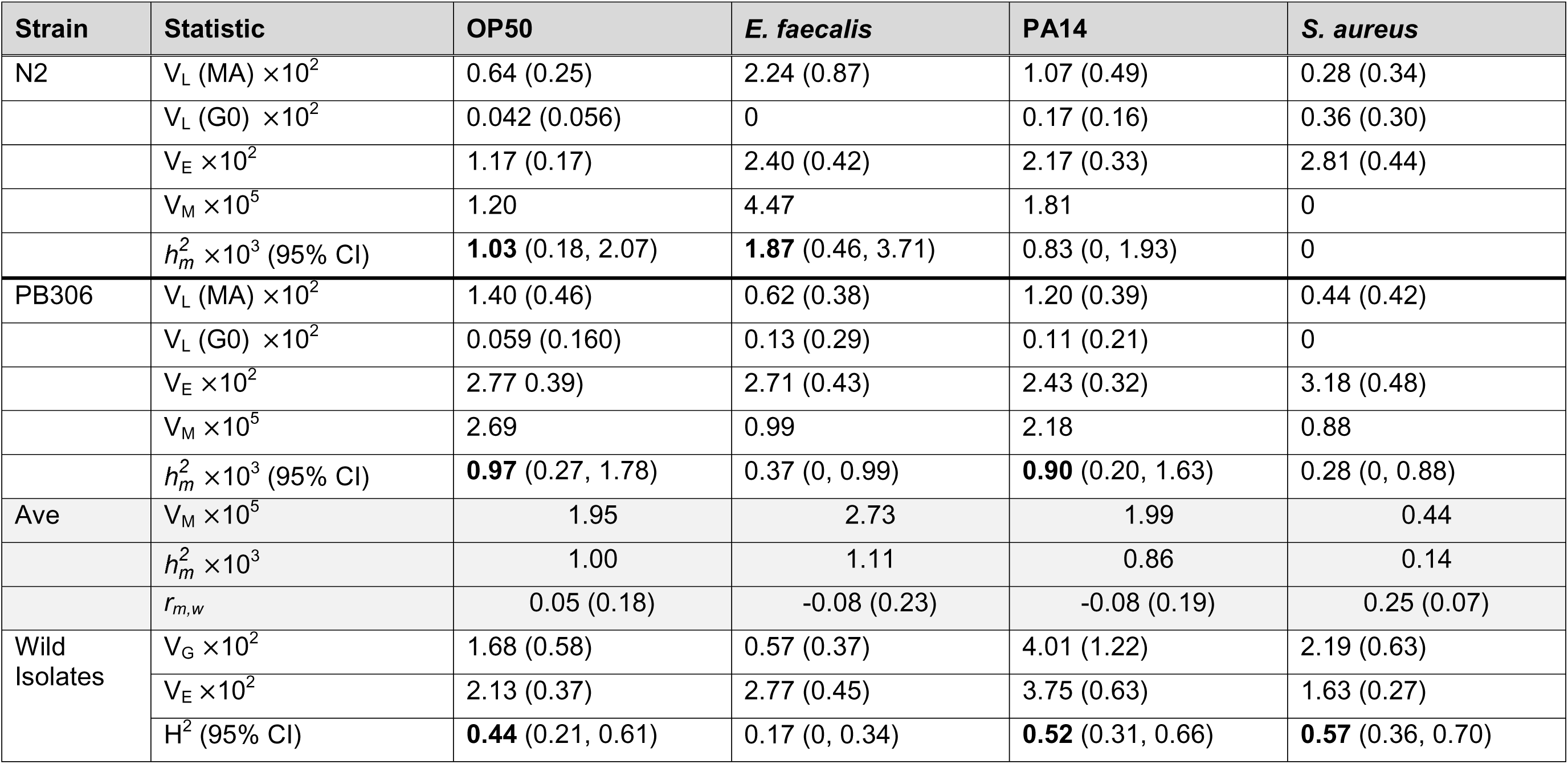
Variances in relative survival, *f_REL_* (SEM). Abbreviations are: V_L_ (among-line variance x 10^2^); V_E_ (environmental variance x 10^2^); V (mutational variance x 10^5^); 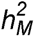(mutational heritability x 10^3^; 95% CI in parentheses); *r_m,w_*(mutational correlation of relative survival with competitive fitness); V_G_ (genetic variance x 10^2^); H^2^ (broad-sense heritability; 95% CI in parentheses). Values of 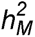 and H^2^ in bold font are statistically significant (p<0.05).

The same analysis for the PS lines has less power than for the MA lines because there are fewer PS lines than MA lines (N2, n=9,9,39,39 for *PA14*, *OP50*, *E. faecalis*, and *S. aureus*, respectively; PB306 n=6,9,39,39). For *E. faecalis* and *S. aureus* there are sufficiently many pseudolines (13/assay block) to credibly test significance of a variance component; in neither strain is V_L,PS_ significantly different from zero for those two bacteria. For OP50 and PA14, we report point estimates of V_L,PS_ and its standard error, and use those estimates in calculations of V_M_, but make no claims about statistical significance of V_L,PS_.

There is significant mutational heritability on two of the four bacterial species in each of the two strains (**Table 2**). Averaged over all bacterial species except *S. aureus*, for which 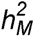 is very low or absent, 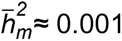, the canonical mutational heritability for a wide variety of traits in many organisms (Houle et al. 1996; Conradsen et al. 2022). We previously estimated mutational parameters for survival on PA14 of (almost) the same set of PB306 MA lines (Etienne et al. 2015). The survival assay (spotted bacterial lawn) and measure of survivorship in that study (LT50) both differed from this study (spread lawn, relative survival at *t_60_*), but the estimated 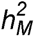 of LT50 was ≈ 0.001, very close to the estimate from this study. The point estimate of 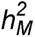 in N2 in this study is nearly the same as that of PB306, although it does not quite meet the 5% threshold of significance. ΔM of PB306 on PA14 was near zero in both studies.

#### (iii) Mutational correlations with competitive fitness

Under the MA conditions, the PB306 MA lines decline in competitive fitness at about 0.1%/generation (ΔM*_w_*= -1.13±0.25x10^-3^/gen), very close to the estimate of ΔM*_w_* for N2 in the same assay (-1.09x10^-3^/gen; Yeh et al. 2018). Pooled over the two sets of MA lines, mutational correlations (*r_M_*) between *f_REL_* on bacterial species *i* and relative competitive fitness are shown in **Table 2**. For all four bacterial species, *r_M_* between *f_REL_* and competitive fitness in the MA environment is ∼0. Evidently mutations (and/or epimutations) that contribute to relative survival have no consistent cumulative effect on competitive fitness under the MA conditions.

### Wild Isolates

There is significant broad-sense heritability (H^2^) for relative survival on three of the four bacterial species, the exception being on *E. faecalis* (**Figure 1C**; **Table 2**). The wild isolate assay did not include pseudoline controls for heritable non-genetic variance, so the estimates of H^2^ are upper bounds on the genetic heritability. Surprisingly, H^2^ is greatest on *S. aureus*, on which there is little if any mutational variance. By way of comparison, we recalculated H^2^ for survival on PA14 from the set of 114 wild isolates reported by Etienne et al. (2015) with LT50 as the summary statistic. Mean LT50 was 63.5 hrs. The recalculated estimate, H^2^=0.41, is slightly less than the estimate from relative survival at *t_60_*in this study (H^2^=0.52).

### Comparison of M and G

One of our primary goals was to compare the mutational architecture of the two sets of MA lines (i.e., **M**_N2_ to **M**_PB306_) to better characterize variation in the mutational process in the context of the multivariate phenotype. We are aware of only one previous such formal comparison, of wing morphology in *Drosophila melanogaster* (Houle and Fierst 2013). However, the lack of significant mutational variance in both sets of lines for two of the four bacterial treatments leads us to think that comparison would not be especially fruitful in this case.

To proceed, we pooled samples over the two sets of MA lines and recalculated among-line (co)variances as outlined in the methods. These (co)variances include the heritable non-genetic component of variance, so we designate the "(epi)mutational" covariance matrix **M\*** to highlight the distinction. Since the assay of the wild isolates was not designed to partition the among-line (co)variance into genetic and non-genetic components (i.e., we estimated **G\*** rather than **G**), **M\*** is the appropriate covariance matrix to compare to **G\***. **M\*** and **G\*** are given in **Tables 3 and 4**, respectively.

**Table 3.** (Epi)mutational (co)variance matrix **M\***. Covariances shown above the diagonal, correlations below the diagonal. Values in bold font are significantly different from 0 (p<0.05). See Methods for details of estimation of **M\***.

| Bacteria | OP50 | *E. faecalis* | PA14 | *S. aureus* |
| --- | --- | --- | --- | --- |
| OP50 | **0.28** (0.08) | 0.04 (0.06) | -0.01 (0.05) | 0.10 (0.05) |
| *E. faecalis* | 0.15 (0.23) | **0.23** (0.09) | 0.10 (0.06) | 0.08 (0.05) |
| PA14 | -0.05 (0.18) | 0.42 (0.22) | **0.25** (0.07) | 0.05 (0.05) |
| *S. aureus* | 0.63 (0.35) | 0.52 (0.41) | 0.36 (0.33) | 0.09 (0.07) |

**Table 4.** (Epi)genetic (co)variance matrix **G\***. Covariances shown above the diagonal, correlations below the diagonal. Values in bold font are significantly different from 0 (p<0.05). See Methods for details of estimation of **G\***.

| Bacteria | OP50 | *E. faecalis* | PA14 | *S. aureus* |
| --- | --- | --- | --- | --- |
| OP50 | **0.31** (0.11) | 0.10 (0.06) | 0.26 (0.12) | -0.05 (0.09) |
| *E. faecalis* | 0.53 (0.31) | 0.11 (0.07) | 0.17 (0.10) | 0.07 (0.06) |
| PA14 | **0.54** (0.20) | 0.60 (0.32) | **0.73** (0.23) | -0.05 (0.12) |
| *S. aureus* | -0.13 (0.24) | 0.31 (0.29) | -0.10 (0.22) | **0.40** (0.12) |

For **M\***, a model with banded main diagonal covariance structure (off-diagonal elements constrained to 0) fit slightly better than a model with unstructured covariance (ΔAICc=1.2; chi-square=11.0, df=6, LRT p>0.08). For **G\***, the unstructured covariance fit slightly better than the model with banded main diagonal (ΔAICc=1.5; chi-square=14.2, df=6, LRT p<0.03). Given the weak support for non-zero off-diagonal elements, it is unsurprising that no consistent pattern emerges from either the mutational or standing genetic correlations, beyond the observation that none of the correlations are large and negative.

The first two principal components explain over 90% of the among-line variance in both **M\*** (60% and 33% for PC1 and PC2) and **G\*** (80% and 17% for PC1 and PC2) (**Supplemental Table S7**). The orientation of **M\*** and **G\*** as determined from the angle θ between **m_max_**and **g_max_** does not differ from the random expectation (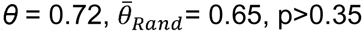; **Figure 2**), from which we conclude that mutation alone is a sufficient explanation for the observed (epi)genetic covariance structure of pathogen susceptibility. The appropriateness of drawing evolutionary inferences from comparisons of principal components of **G** has been criticized on the grounds that "…(evolutionarily) causal factors need not have orthogonal effects on the phenotype" (Houle et al. 2002), but when most of the heritable variance falls along PC1, the "genetic lines of least resistance" approach seems reasonable.

**Figure 2.**
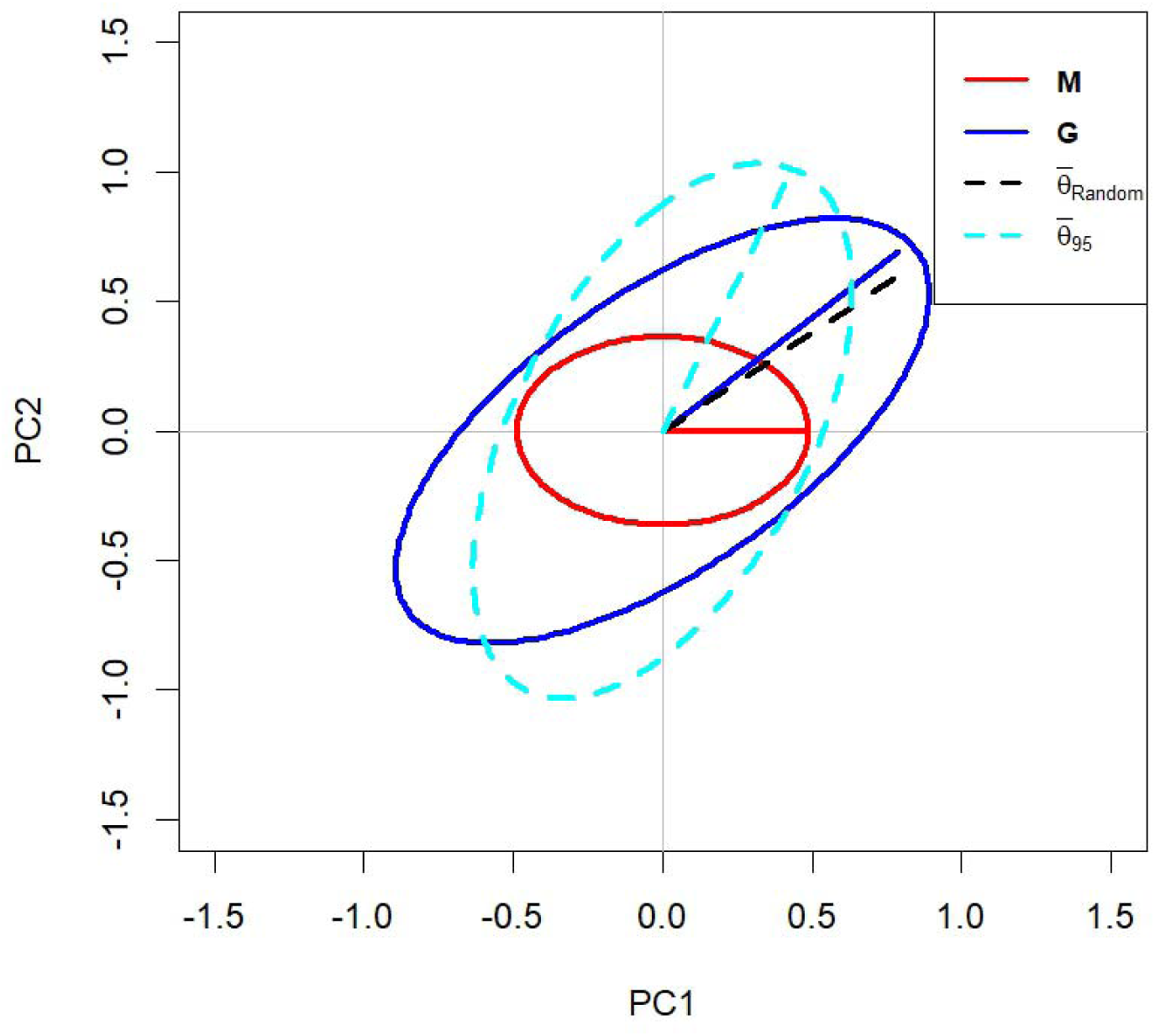
Relative orientation of the first two principal components of the mutational covariance matrix **M\*** (red ellipse) and the standing genetic covariance matrix **G\*** (blue ellipse). Solid colored lines represent PC1 (**m_max_** and **g_max_**, respectively). Dashed black line depicts the mean difference between random pseudoMA lines and pseudoWild Isoates (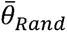; see Methods for explanation). Light blue dashed ellipse shows the 95% confidence limit of the deviation between pseudoMA lines and pseudoWild Isolates.

### The Strength and Form of Natural Selection

The finding that ΔM<0 in every case implies that mutations that reduce survival upon exposure to pathogenic bacteria are deleterious, on average. That is not surprising in hindsight, although it is certainly plausible that fitness tradeoffs could result in stabilizing selection on pathogen susceptibility. The lack of correlation between survival on pathogens with competitive fitness in the MA environment indicates that pleiotropy between mutational effects in the two contexts is not strong, but reveals little about selection in the natural environment.

To get a rough idea of how strong selection on these traits is, the neutral expectation provides a benchmark. For a neutral trait in a predominantly selfing organism such as *C. elegans*, at mutation-drift equilibrium, V_G_≈4*N_e_*V_M_ (Lynch and Hill 1986). *N_e_* in *C. elegans* has been estimated to be on the order of 10^4^-10^5^ (Gilbert et al. 2021), so if V_M_≈2x10^-5^ (the average for relative survival on OP50), we expect V_G_≈0.8-8, about 40-400 times greater than the observed value of ∼0.02. The discrepancy is similar for PA14, and slightly greater for *E. faecalis*. Although there is considerable uncertainty associated with all of these estimates, the ability to survive exposure to those three species of bacteria is clearly not a neutral trait; mutations affecting survival on those bacteria are subject to effective purifying selection. *S. aureus* may present a different scenario; we return to it below.

If selection is sufficiently strong to be approximately deterministic (i.e., ignoring drift), at mutation-selection balance, 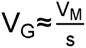, where *s* represents the average strength of selection against a mutation affecting the trait (Barton 1990); see Charlesworth (2015) for caveats. By that calculation, *s* ≈ 0.001 for mutations affecting relative survival on OP50, roughly the same on the virulent pathogen PA14 and a bit stronger on *E. faecalis*. By way of comparison, a similar calculation for competitive fitness (measured in the MA conditions) revealed *s*≈0.005 (Yeh et al. 2018; the value is halved if V_G_=V_L_ rather than V_L_/2), which is in close agreement with a direct estimate of the average mutational effect 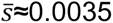, calculated by dividing the cumulative decline in competitive fitness of the MA lines by the average number of mutations carried by an MA line. These point estimates are obviously rough, but it is evident that mutations that affect susceptibility to pathogens experience significant selection in nature. Whether the selection is direct, imposed by exposure to pathogens in nature, or indirect, due to pleiotropic effects on fitness in the absence of pathogens, cannot be ascertained from these experiments. However, the absence of a mutational correlation between relative survival and relative fitness (in the MA context) implies that it might be the former. Recent studies have revealed a complex transgenerational epigenetically inherited (TEI) behavioral response specific to the PA14 strain of *P. aeruginosa* that varies among wild isolates (Moore et al. 2021), which reinforces the inference that PA14 imposes direct selection in nature.

Mutation-selection balance (MSB) is not the only mechanism by which genetic variation may be maintained; balancing selection (BS) is another. There are several well-documented balanced polymorphisms in *C. elegans* (summarized in Lee et al. (2021)), and there are numerous examples from diverse organisms in which pathogens do contribute to BS (Fumagalli et al. 2009; Hawley and Fleischer 2012; Horger et al. 2012; Quéméré et al. 2021). If BS is the predominant mechanism responsible for the maintenance of genetic variation, there is no reason to expect V_G_ and V_M_ to be correlated. In fact, V_M_ and V_G_ are nearly always highly positively correlated (Houle 1998; McGuigan et al. 2015; Farhadifar et al. 2016). BS and MSB are often set up as alternative hypotheses for the maintenance of genetic variation, but it is realistic to expect that uniformly deleterious alleles will always contribute to trait variation, whereas balanced polymorphisms may or may not. If that scenario is generally true, the effect of BS would be to weaken the relationship between V_M_ and V_G_, and that does not seem to be the case. If, in contrast, BS is the exception rather than the rule, the occasional trait experiencing BS would appear as a high outlier, with more V_G_ than predicted by its V_M_, and selection as inferred from the ratio of V_M_ to V_G_ would appear anomalously weak.

Relative survival on *S. aureus* seems to fit that pattern, at least at first glance. Averaged over both sets of MA lines, V_M_ for relative survival on *S. aureus* is only about 20% of V_M_ on the other bacteria, whereas V_G_ is not atypically low, and H^2^ is the highest of the four bacteria. If *N_e_*is closer to 10^4^ than 10^5^, the selection coefficient inferred from V_M_/V_G_ (*S* ≈ 2 x 10^-4^) is approaching effective neutrality (1/*N_e_*, the "drift barrier"). However, averaged over the two sets of MA lines, ΔM is the greatest (most negative) on *S. aureus*, which implies that accumulated (epi)mutations significantly reduce relative survival on *S. aureus*.

It is not unheard of in the MA literature for trait means to change significantly even in the absence of significant mutational variance (Shabalina et al. 1997 is a well-cited example in Drosophila). It is a recurring phenomenon in *C. elegans* MA experiments, and it is most common for traits that have inherently high variability (e.g., male competitive mating success, Yeh et al. 2018; metabolite concentration, Johnson et al. 2020). The phenomenon may be more detectable in *C. elegans* than in other systems, for two related reasons. The ability to easily maintain cryopreserved stocks permits the contemporaneous assay of the unevolved G0 ancestor and the MA lines, and transgenerational epigenetic inheritance (TEI) is extremely well-characterized in *C. elegans* (Rechavi and Lev 2017; Beltran et al. 2020). If TEI contributes to the heritable variance in an unbiased way and of similar magnitude to mutational variance (Johnson et al. 2020), it could overwhelm the signal of mutational variance even in the presence of a directional effect of mutations. Alternatively, if a trait is especially sensitive to microenvironmental noise, V_E_ may overwhelm V_M_ such that among-line variance is undetectable at the given sample sizes. That may be the case for *S. aureus* in the MA lines, for which V_E_ is greatest on *S. aureus* in both the N2 and PB306 lines. However, to make things even less clear, V_E_ is smallest on *S. aureus* in the wild isolates.

Given the time- and labor-intensive properties of both MA experiments and pathogen survival assays, it is perhaps unsurprising that there are very few studies to have quantified the effects of spontaneous mutations on pathogen susceptibility. Recently, Mendel et al. (2024) investigated the effects of spontaneous mutations on the susceptibility of *Drosophila serrata* to Drosophila C virus (DCV). They reported a mutational heritability for LT50 of 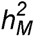 ≈ 0.004-0.005, significantly greater than the estimates of 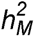 in this study and toward the high end of estimates of 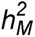 The mutational properties (rate and spectrum) at the genomic level in *D. serrata* remain unpublished, but recent estimates from parent-offspring sequencing of *D. melanogaster* and *D. simulans* revealed a base substitution/small indel rate 1-2X that of *C. elegans* (Wang et al. 2023). The genomes of those two fly species are somewhere between 36% (*D. simulans*) and 67% (*D. melanogaster*) larger than that of *C. elegans*, so the mutational target is probably somewhat larger. However, the per-genome, per-generation rate of transposable element insertions is about 50% greater than that of new base-substitutions/small indels. Based on limited data, we infer that spontaneous transpositions constitute < 1% of all spontaneous mutations (Saxena and Baer, unpublished results), so if *D. serrata* has similar TE activity to the other Drosophila species, that could explain the greater mutational heritability compared to *C. elegans*. Of course, there are many other differences between flies and worms, and between bacteria and viruses. Somewhat surprisingly, *C. elegans* is only known to have one viral pathogen (the Orsay virus, Felix et al. 2011), which has been established as a laboratory model system (Batachari et al. 2024). The MA lines and wild isolates are freely available and await the intrepid Ph.D student to do the investigation.

## Conclusions

1. The cumulative effect of mutations is to increase susceptibility to pathogens. Moreover, survival upon exposure to pathogens is uncorrelated with competitive fitness under the MA conditions. Thus, there is no evidence of tradeoffs between susceptibility to pathogens and fitness in the absence of pathogens.
2. To the extent that **M\*** and **G\*** are reliable estimators of **M** and **G**, respectively, it is evident that survival on these bacteria is subject to effective selection (because **G**<< 2*N_e_***M**), but also that **M\*** is a sufficient predictor of the orientation of **G\*** in multivariate trait space.
3. The number of MA lines included (124), and the number of generations of MA (250, ∼1000 days) are quite large. The number of wild isolates (47) is not so large, but is not small. Nevertheless, the results are (depending on one’s perspective) either intriguingly or distressingly variable between strains (N2, PB306) and bacteria. Nevertheless, the one point of direct comparison – PB306 on PA14 – yielded results that are very consistent with a previous study. In our opinion, the way forward lies in development and adoption of high-throughput phenotyping methods to increase the practical sample sizes in these kinds of assays.

## Data Availability

All data are included as Supplemental Tables and will be deposited in Figshare upon acceptance for publication. All code is available in Figshare at 10.6084/m9.figshare.28046366.

## Acknowledgments

Q. Allen, J. Dembek, D. Feistel, M. Puentes, A. Shoucair, and M. Snyder provided assistance in the laboratory. Support was provided by NIH awards GM107227 and GM127433.

