## Supplemental Material for "Cumulative effects of mutation and selection on susceptibility to bacterial pathogens in *Caenorhabditis elegans*"

**1. Slow Killing Assay (SKA).**

Preparation of slow-killing assay (SKA) plates – Bacterial strains were acquired from Erik Andersen (Northwestern University) and kept frozen at -80°C until use. On day 5 of the assay, the bacterial species to be used in that block was thawed and spread onto LB plates and incubated at 37°C for two days. On day 7, a single bacterial colony was picked to inoculate ~28 ml of YT media, which was incubated in a shaking incubator at 100 RPM at 37°C for 24 hrs. After 24 hours (day 8), 5μl of bacterial culture was spread onto a 35 mm NGMA SKA plate containing 0.05% of 100mg/mL filter-sterilized fluorodeoxyuridine (FUDR) which prevents worm progeny from hatching by inhibiting DNA synthesis (Hosono 1978). Assay plates were incubated in a closed plastic container at 37°C for 24 hours. On day 9, SKA plates were moved to a 25°C incubator.

Assay protocol - On day 1 of an assay block the MA lines and G0 ancestor were thawed onto 60mm NGMA agar plates (P0) seeded with OP50 and stored at 20°C. After 24 hours (day 2) a single L4 hermaphrodite (P1) was picked onto a new 60mm agar plate seeded with OP50 and incubated at the ultimate assay temperature of 25°C. Each MA line was replicated three-fold; the number of replicates from PS lines varied by block (**Supplemental Table S4**). On day 6, a single L4 hermaphrodite (F1) from each replicate was picked to a new plate and incubated at 25°C. On day 10, 30±3 L4 (sometimes L3) stage worms (F2) from each line were picked onto the SKA plates. Assay plates were assigned random numbers and subsequent handling was in random order. The number of living progeny on each plate was counted at 12 hour intervals for 120 hours.

Survival was determined via blue light exposure. Worms exhibit a phototactic response when exposed to blue light (Lee and Aschner 2016). Non-motile worms were illuminated with a blue laser pointer for five seconds; worms that did not move after this time were categorized as dead. It is important to note that only worms that were observed moving were counted as alive, those that may have crawled off the plate, or buried themselves into the agar were not counted as alive.

**2. Estimation of mutational and standing genetic covariance matrices (M and G).**

The diagonal elements of the covariance matrix are the variances in *f_RE_*_L_ for bacterial species *i* and the off-diagonal elements are the covariances between *f_REL_* for bacteria *i* and *j*. (Co)variances can be partitioned into an among-line and within-line (residual) component. To estimate **M**, we pooled the data from the two strains, inasmuch as no one strain can lay claim to "the" mutational architecture of its species. For each treatment (MA or G0), the full GLM is *y'_ijk_=μ+L_j|i_+e_jk|i_*, where *y'_ijk_* is the variance-standardized (Studentized) residual of *f_REL_* of a replicate from the assay block mean, *μ* is the overall mean, *L_j|i_* is the random effect of line *j* (MA or PS) on bacteria *i*, and *e_k|ij_* is the residual effect. The *L_j|i_* are distributed (approximately) ~N($\bar{\text{a}}$,***G***), with mean $\bar{\text{a}}$ equal to the treatment mean and covariance structure ***G***, as described below. The *e_k|ij_* are assumed normally distributed ~N(0,***R***) with mean 0 and covariance structure ***R***, where the off-diagonal elements of ***R*** are constrained to equal 0 (banded main diagonal covariance), since the replicates assayed in the different pathogens are biologically independent samples.

For each treatment (MA or G0), we tested two (G0) or three (MA) hierarchical models of covariance structure, (1) the full model, with unstructured ***G***, (2) banded main diagonal ***G***, in which the off-diagonal elements are constrained to 0, and (3) ***G***=**0**, in which all elements of ***G*** are constrained to 0. The best model was decided based on the corrected AIC (Hurvich and Tsai 1989). If the best model had more parameters than the next-best model, the models were compared by likelihood-ratio test (LRT); the models are nested, so twice the difference in the log(likelihood) is asymptotically chi-square distributed with degrees of freedom equal to the difference in the number of parameters in the two models. For the G0 PS lines, ***G*** is necessarily banded main diagonal (if not **0**), because the PS lines are independent biological samples in the different pathogen assays.

The genetic covariance matrix of the wild isolates, **G**, was estimated similarly to **M**. Relative survival was standardized by the mean of the N2 G0 pseudolines on OP50 (see main text for explanation). The GLM is the same as for the MA lines, where "Line" refers to the wild isolate. Covariances were estimated jointly from the unconstrained model and compared to the model constrained to banded main diagonal structure. Note the notational distinction between ***G*** (italicized) and **G** (no italics); the former represents the estimated among-line variance-covariance matrix of group *i*, and the latter represents the genetic variance-covariance matrix.

**3. Competitive fitness assay.**

Competitive fitness of hermaphrodites was assayed in two blocks beginning in May, 2005. The cryopreserved G0 ancestor of the MA lines was thawed and 20 replicate populations initiated from a single L3/L4 stage worm placed on a standard 60 mm NGM agar plate seeded with 100 µl of an overnight culture of the OP50 strain of *E. coli*. These populations are referred to as "pseudolines" and designated the P0 generation. Seven L3/L4 stage offspring from each pseudoline were transferred singly to new plates, designated the P1 generation. Pseudolines were subsequently treated identically to MA lines. G250 MA lines were thawed and seven revived L3/L4 stage worms from each line were placed individually on standard 60 mm NGM plates, labeled P1. All P1 plates were assigned a unique random number and all subsequent experimental manipulations were performed in sequence by random number. Replicates were maintained for two more generations (P2-P3) by transfer of a single L3/L4 stage offspring at four-day intervals to control for parental and grandparental effects. At the same time, we thawed a replicate of the GFP-marked competitor strain ST2 and made several large replicate populations by transferring a chunk from the initial plate to a new 10 cm plate.

On the second day after the P3 worm began reproduction, a competition plate for each replicate was set up by transferring a single L1-stage larva from the P3 plate and a single L1-stage ST2 competitor onto a 60 mm NGM agar plate seeded with 100 µl of the HB101 strain of *E. coli* and supplemented with nystatin to retard fungal contamination. Competition plates were incubated at 20° C for eight days, at which point food was exhausted. Worms were washed from competition plates in cold M9 buffer, settled on ice and 100 µl of the settled worms transferred into a drop of glycerol on the lid of an empty 60 mm agar dish and the bottom of the empty dish pressed into the lid. The glycerol immobilizes the worms and pressing them between halves of the plate puts them into the same focal plane. We took two pictures of each plate at 40X magnification through a Leica MZ75 dissecting microscope fitted with a 100 W mercury arc lamp and epifluorescence GFP filter cube (470/40 nm excitation filter, 525/50 nm emission filter) using a Leica DFC280 camera connected to a computer running the Leica IM50 software (Leica Microsystems Imaging Solutions Ltd). The first picture used the arc lamp and GFP filter cube (the "green" image) and the second, taken immediately afterwards, used transmitted white light (the "white" image). All worms are visible in the white image, whereas wild-type (non-GFP) worms appear only faintly in the green image. The difference between the number of worms in a white image and in the matching green image is the number of focal worms in the sample.

Images were imported into ImageJ software (<http://rsb.info.nih.gov/ij/>) and worms were counted as follows. If there appeared to be fewer than 200 worms visible in a white image, we first counted every worm in the white image and then each worm visible in the accompanying green image. If there appeared to be > 200 worms in the white image we drew a rectangle around approximately 200 worms and counted them. We then pasted the same rectangle in the green image and counted the worms visible within the rectangle.

*Data Analysis* -

i) Measures of competitive fitness - Competitive fitness has two components: (1) did the focal individual reproduce at all? If not, relative fitness is zero regardless of the number of offspring of the competitor, and (2) given that the focal individual did reproduce, what fraction of the offspring belong to the focal individual? Given that a focal individual did reproduce, the ratio *p/(1-p)* is related to competitive fitness by the relationship

$\frac{p_{t}}{q_{t}}=\frac{p_{0}}{q_{0}}\left( \frac{W_{foc}}{W_{C}} \right)^{t}$	Equation 1

(Barton et al. 2007, equation 17.2), where *t* represents the number of generations in the fitness assay, *p_0_* is the frequency of the focal type (G0 or control) at the beginning of the assay, *p_t_* is the frequency of the focal type (G0 control or MA) at the conclusion of the assay, *q* = 1-*p*, *W_foc_* is the absolute fitness of the focal type, and *W_C_* is the absolute fitness of the competitor. Each trial was started with one focal worm and one competitor, so the ratio $\frac{p_{0}}{q_{0}}=1$. We refer to the ratio *p*/(1-*p*) as the "competitive index", *CI* (Shabalina et al. 1997). *CI* provides a measure of fitness of the focal type relative to the competitor, raised to the power *t*. All analyses of *CI* were performed on natural log-transformed data.

ii) Probability of reproduction, *π* - Probability of reproduction is a binary trait. If a focal worm reproduced the replicate is scored as a success ("event=1"); if the focal worm did not reproduce it is scored as a failure ("event=0"). Data were analyzed by Generalized Linear Mixed Model (GLMM) with estimation by Residual Subject-specific Pseudolikelihood (RSPL) as implemented in the GLIMMIX procedure of SAS v.9.4 with a logit link function and a random residual. Treatment (MA vs. Control) is a fixed effect and Line and Replicate (nested within Line) are random effects. Block is a random effect in principle. However, pseudolikelihoods are not appropriate criteria for model selection (e.g., by AIC; SAS/Stat User's Guide (SAS Institute 2013)), so rather than include or exclude variance components including block on the basis of estimates for which there is little power (because *n*=2), we chose to model block as a fixed effect for this analysis. It is common in the analysis of MA fitness assays to treat block as a fixed effect when the number of blocks is small (e.g., Houle et al. 1994; Shaw et al. 2000).

Each line (MA and G0 pseudoline) was assayed for probability of reproducing, *π*. The full model is written as:

*π_ijkl_ = µ + t_k_ + b_j_ + l_l|jk_ + ε_il|jk_*

where *π_ijkl_* is a binary variate scored as 1 if the focal worm produced at least two offspring and 0 if it did not, *µ* is the overall mean, *t_k_* is the fixed effect of treatment *k* (G0 or MA), *b_j_* is the fixed effect of block *j*, *l_l|jk_* is the random effect of line (or pseudoline) *l*, conditioned on block and treatment, and *ε_il|jk_* is the random residual, conditioned on block and treatment. Random effects were estimated separately for each block/treatment combination by means of the GROUP option in the RANDOM statement of the GLIMMIX procedure. Significance of fixed effects was determined by F-test of Type III sums of squares. The distribution of *π* was strongly left-skewed, so means and standard errors were calculated by an empirical bootstrap procedure (Efron and Tibshirani 1993; Baer et al. 2006). Resampled datasets were constructed by resampling lines within blocks, followed by estimation of means and variance components from the GLMM described above.

Competitive Index (*CI*) – *C*I was analyzed using a standard general linear model (GLM) as implemented in the MIXED procedure of SAS v. 9.4. Studentized residuals of natural log-transformed data were scrutinized for outliers by eye against a Q-Q plot. After removal of three outliers (*n* = 671), the data were initially fit to the linear model:

*y_ijkl_ = µ + t_k_ + b_j|k_ + c_jk_ + l_l|jk_ + ε_il|jk_*

where *y_ijkl_* is the log(*CI*) of the individual replicate and the independent variables are defined as in the previous section. Block was modelled as a random effect in this analysis. Variance components of random effects were estimated by restricted maximum likelihood (REML). The among-block component of variance was estimated separately for each treatment and the among-line and among-replicate (nested within line) components were estimated separately for each treatment/block combination by means of the GROUP option in the RANDOM or REPEATED statement of the MIXED procedure (Fry 2004).

We first analyzed the full model above, then sequentially simplified the model by first pooling the random effects across grouping levels (e.g., estimating a single among-line variance rather than estimating it separately for each block) and then removing the effect entirely. The model with the smallest corrected AIC (AICc) was chosen as the best model, and significance of the fixed effect of treatment (MA or G0) in that model was determined by F-test of Type III sums of squares, with degrees of freedom determined by the Kenward-Roger method (Kenward and Roger 1997). If two models had equal AICc, the simpler model was chosen as the best model. In addition, we calculated empirical bootstrap estimates of the mean and standard error of *CI_M_*, resampling over lines within blocks followed by estimation of means and variance components from the GLM described above.
