## Supplemental Figures S1-S5 for "Cumulative effects of mutation and selection on susceptibility to bacterial pathogens in *Caenorhabditis elegans*"

s

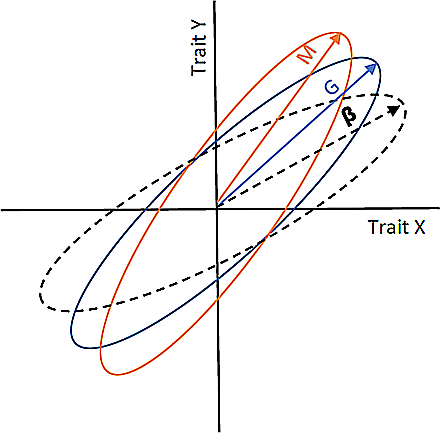

**Supplemental Figure S1**. Depiction of a hypothetical relationship between two traits, X and Y. The orange ellipse represents the input of genetic (co)variance by mutation (**M**), the black dashed ellipse represents the trajectory of bivariate directional selection (**β**), and the blue ellipse represents the genetic (co)variance matrix (**G**) at mutation-selection balance. After [16].

a

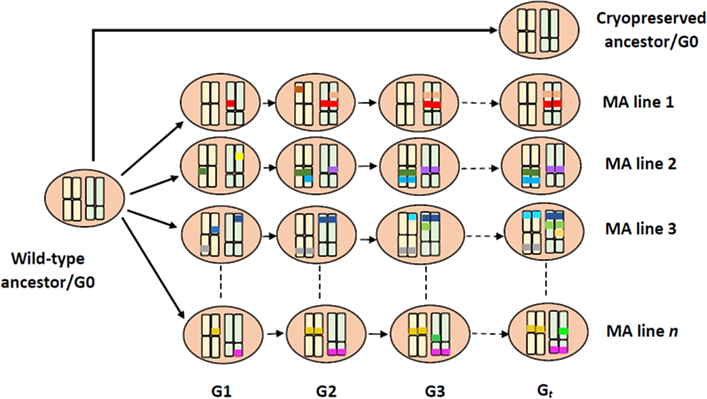

**Supplemental Figure S2**. Schematic depiction of the MA experiment. Ovals represent individuals, with two pairs of chromosomes (yellow, green). Colored bars on chromosomes represent mutations unique to each MA line, which accumulate over the course of the t (≈250) generations of MA; the ancestor is taken to be genetically uniform and homozygous at all loci. At each generation (G1, G2, ...G_t_), a single individual is propagated to a new plate to found the next generation.

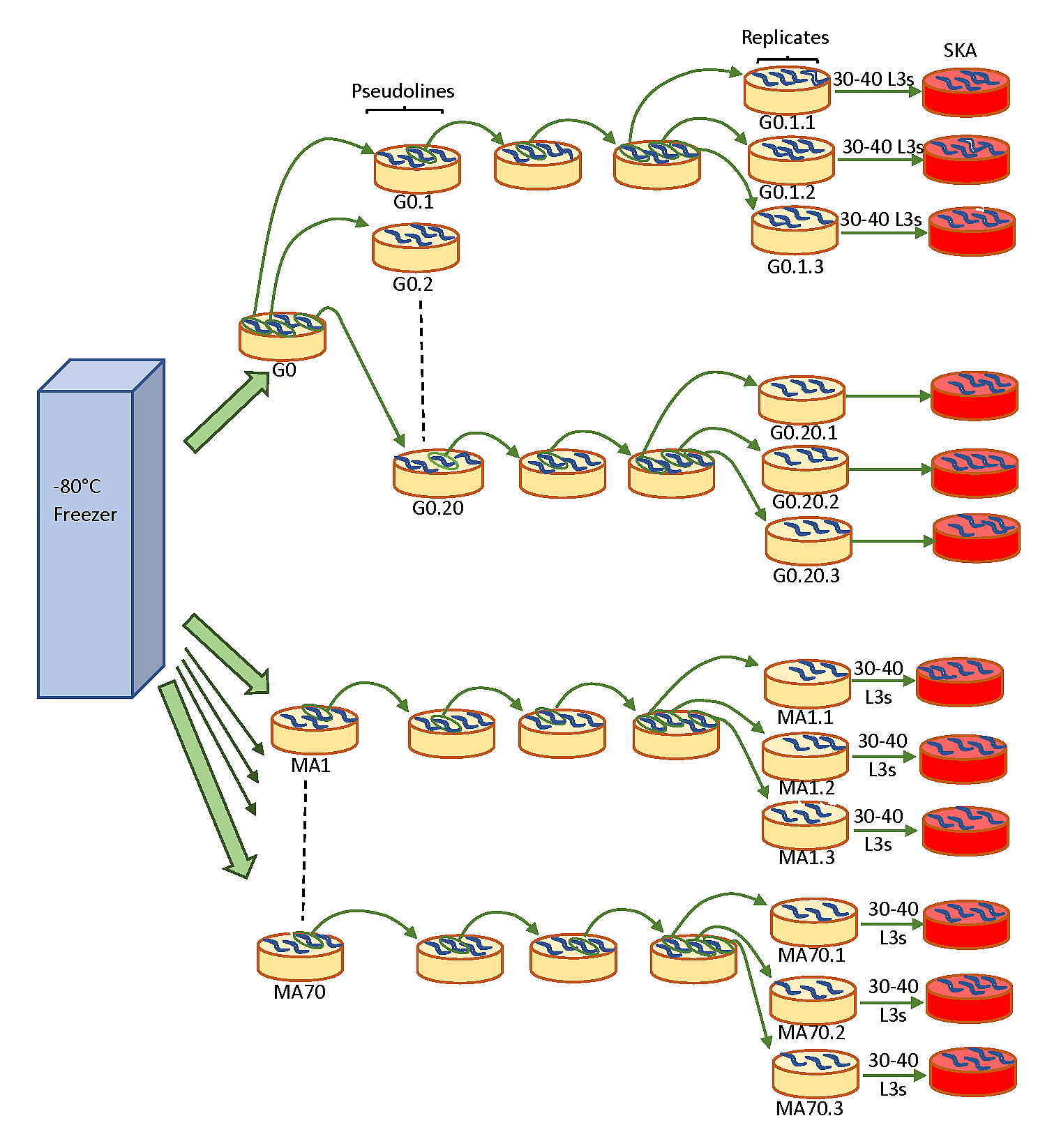

**Supplemental Figure S3**. Schematic depiction of the Slow Killing Assay (SKA). See Methods for details of the assay.

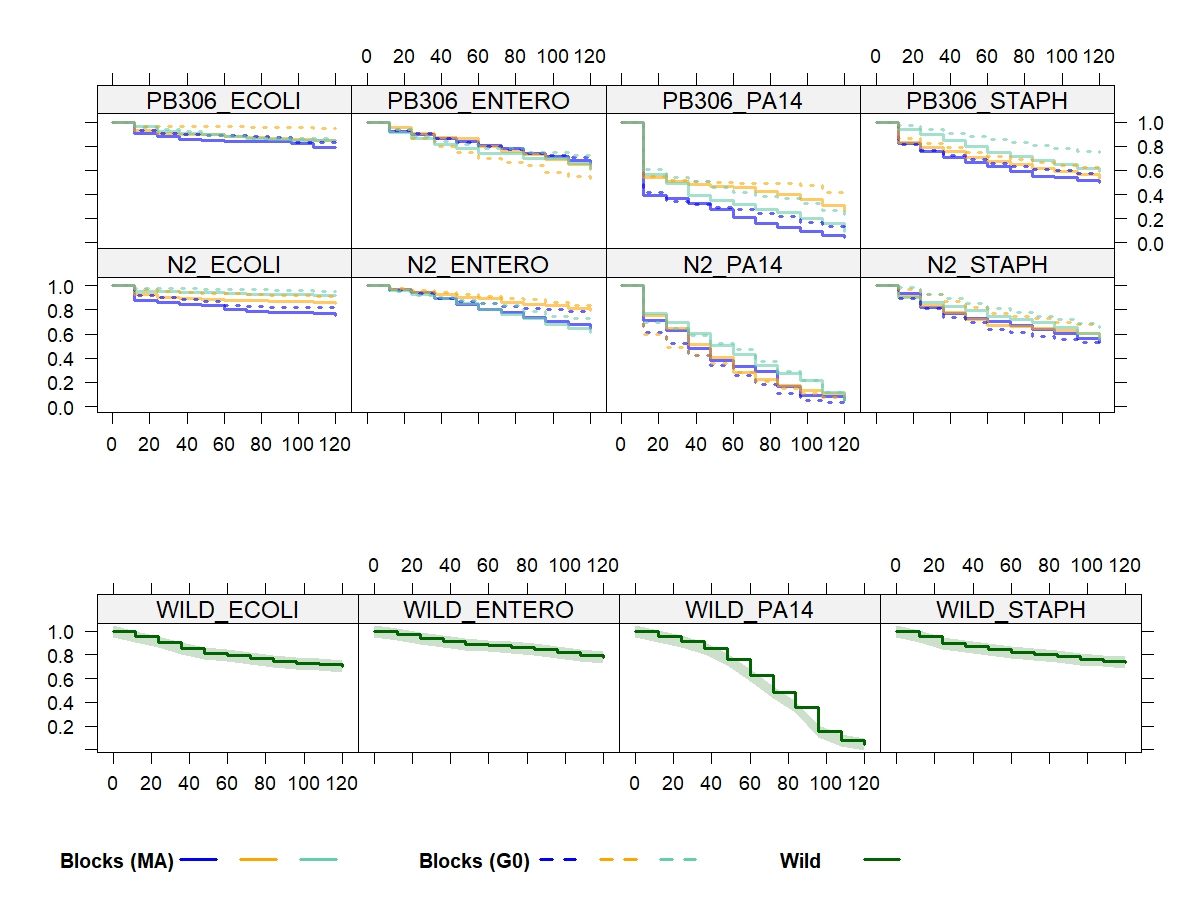

**Supplemental Figure S4.** Survivorship curves (block means) for MA/PS lines on the four bacterial species. Top panels, N2; bottom panels, PB306. Left to right (**A-D**): (**A**) OP50 (ECOLI); (**B**) E. faecalis (ENTER); (**C**) PA14; (**D**) S. aureus (STAPH). Solid lines designate MA lines, dashed lines designate G0 pseudoline (PS) means.

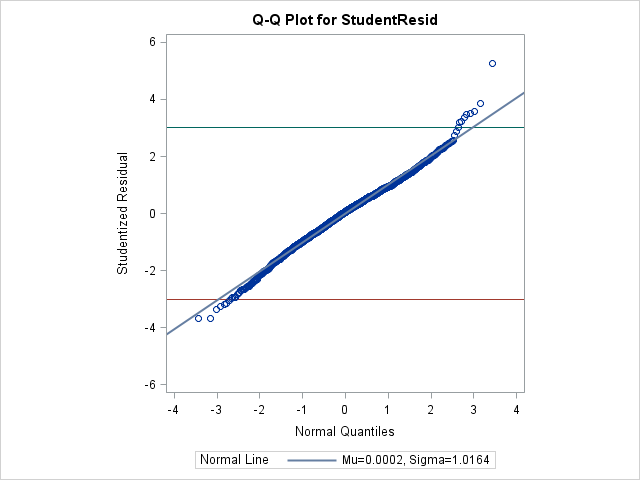

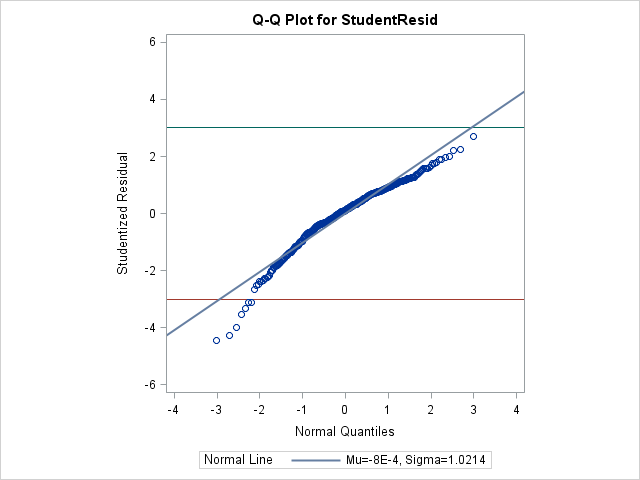

**Supplemental Figure S5**. Q-Q plots of studentized residuals of relative survival (*f_REL_*) from the linear model described in the Methods. (A) Top panel, all MA and G0 lines, pooled over strains, treatments, pathogens, and assay blocks. (B) Bottom panel, wild isolates, pooled over pathogens. Samples with absolute studentized residual +/-3 were removed from subsequent analyses.
